# Robust Preclinical Evidence in Somatic Cell Genome Editing: A Key Driver of Responsible and Efficient Therapeutic Innovations

**DOI:** 10.1101/2020.09.09.290338

**Authors:** Merlin Bittlinger, Johannes Schwietering, Daniel Strech

**Affiliations:** Charité – Universitätsmedizin Berlin, corporate member of Freie Universität Berlin, Humboldt-Universität zu Berlin, and Berlin Institute of Health Germany; QUEST – Center for Transforming Biomedical Research, Berlin Institute of Health Germany (BIH) Anna-Louisa-Karsch-Str. 2, 10178 Berlin, Germany; Hannover Medical School, Institute for Ethics, History, and Philosophy of Medicine, Hannover, Germany

## Abstract

Somatic cell genome editing (SCGE) is highly promising for therapeutic innovation. Multifold financial and academic incentives exist for the quickest possible translation from preclinical to clinical studies. This study demonstrates that the majority of 46 preclinical SCGE studies discussed in expert reviews as particularly promising for clinical translation do not report on seven key elements for robust and confirmatory research practices: (1) randomization, (2) blinding, (3) sample size calculation, (4) data handling, (5) pre-registration, (6) multi-centric study design, and (7) independent confirmation. Against the background of the high incentives for clinical translation and recent concerns about the reproducibility of published preclinical evidence, we present the here examined reporting standards (1-4) and the new NIH funding criteria for SCGE research (6-7) as a viable solution to protect this promising field from backlashes. We argue that the implementation of the novel methodological standards, e.g. “confirmation” and “pre-registration”, is promising for preclinical SCGE research and provides an opportunity to become a lighthouse example for trustworthy and useful translational research.

## Background

The field of somatic cell genome editing is rapidly evolving with recent major innovations.^1,2^ There is a great potential for therapeutic breakthroughs in many important medical conditions. In 2010, zinc finger nucleases (ZNF) emerged as the first gene editing platform to translate into clinical trials.^3^ In 2015, the genome editing platform TALEN has been applied for the first time in humans in the context of “compassionate use” in pediatric patients under regulation of UK Medicines and Healthcare products Regulatory Agency (MHRA).^4,5^ In 2016, CRISPR\Cas9 was applied for the first time in humans.^6^ In 2018, the U.S. Food and Drug Administration (FDA) granted permission for the first Investigational New Drug (IND) application (EDIT-101) for non-heritable *in vivo* genome-editing.^7^ Quickly thereafter the FDA granted also an IND Fast Track Designation (CTX001).^8^ Current clinical trials span diverse therapeutic areas such as clinical trials for Duchenne muscular dystrophy, human immunodeficiency virus (HIV) infection, hemophilia B, acute myeloid leukemia, among several others.^9^

Due to the high therapeutic promise, there are high incentives for translational research leveraging genome editing technologies. Moreover, there are also high economic expectations and enormous financial investments with private and public research funding. The roughly estimated net worth of biotechnology companies specialized on genome editing ranges from less than US$ 4 billion (Cellectis,^10^ Sangamo,^11^ Intellia,^12^ or CRISPR Therapeutics^13^) to up to US$ 57 Billion (Vertex Pharmaceuticals Inc.). In this rapidly evolving and highly competitive field “the race to the market needs a rapid turnaround of data”.^14^

With all these expectations and incentives in place, we argue that the trustworthiness and usefulness of preclinical evidence is of utmost importance for quickly translating preclinical genome editing research into clinical trials. Trustworthy and useful evidence requires, first of all, the employment of measures to reduce validity threats.^15,16^ According to results of a multi-stakeholder workshop at the NIH in 2012 (so-called “Landis-4” criteria)^17^ and the revised submission guidelines of Nature journals,^18^ the measures to increase scientific validity include at minimum the use of (1) randomized group allocation, (2) blinded outcome assessment, (3) a sample-size calculation, and (4) the transparent explanation of exclusion of animals or data from analyses.^17^ Further yet less established measures in preclinical research that would reduce risk of bias and increase reproducibility of published preclinical evidence are (5) the public pre-registration of study protocols,^19^ (6) performing multi-centric studies involving primary data collection at two or more laboratory sites,^20,21^ and an (7) independent validation to “confirm” promising preclinical results before translation.^22,23^

The objective of this scoping review is to understand how well the field of preclinical genome-editing research is already meeting the well-established reporting standards for internal validity (1-4). A second objective is to explore to what degree the additional measures to increase the reproducibility and robustness of preclinical findings (5-7) are already implemented in the field of somatic genome editing.

## Methods and Materials

To identify preclinical studies that promote the clinical translation of a medical intervention using somatic genome editing, we searched on PubMed/MEDLINE for relevant review articles about the state of clinical translation in the field of somatic cell genome editing. The search string was adopted from a recent systematic review of preclinical studies^24^ and informed by preceding systematic reviews (Figure 1).^25^ We assessed the review articles for mention of preclinical studies. Criteria for inclusion into analysis were, first of all, the reporting of (a) at least one animal experiment, which involves (b) either CRISPR/Cas, TALEN, or ZFN-based gene editing machinery as intervention. Second, included studies needed (c) to report an outcome-of-interest that is explicitly related to clinical application. Exclusion criteria were (i) duplication of articles in many different reviews, (ii) development of new animal models for human diseases, (iii) germline modification as intervention, (iv) *in vitro* experiments, or (v) pure basic science. If studies met these inclusion criteria, we assessed the reporting of the design aspects (1-7) mentioned above. During the piloting of the search string, five candidate studies were identified and also included into the eligibility assessment based on the full text article (see https://osf.io/e6zqh/).

**Figure 1.**
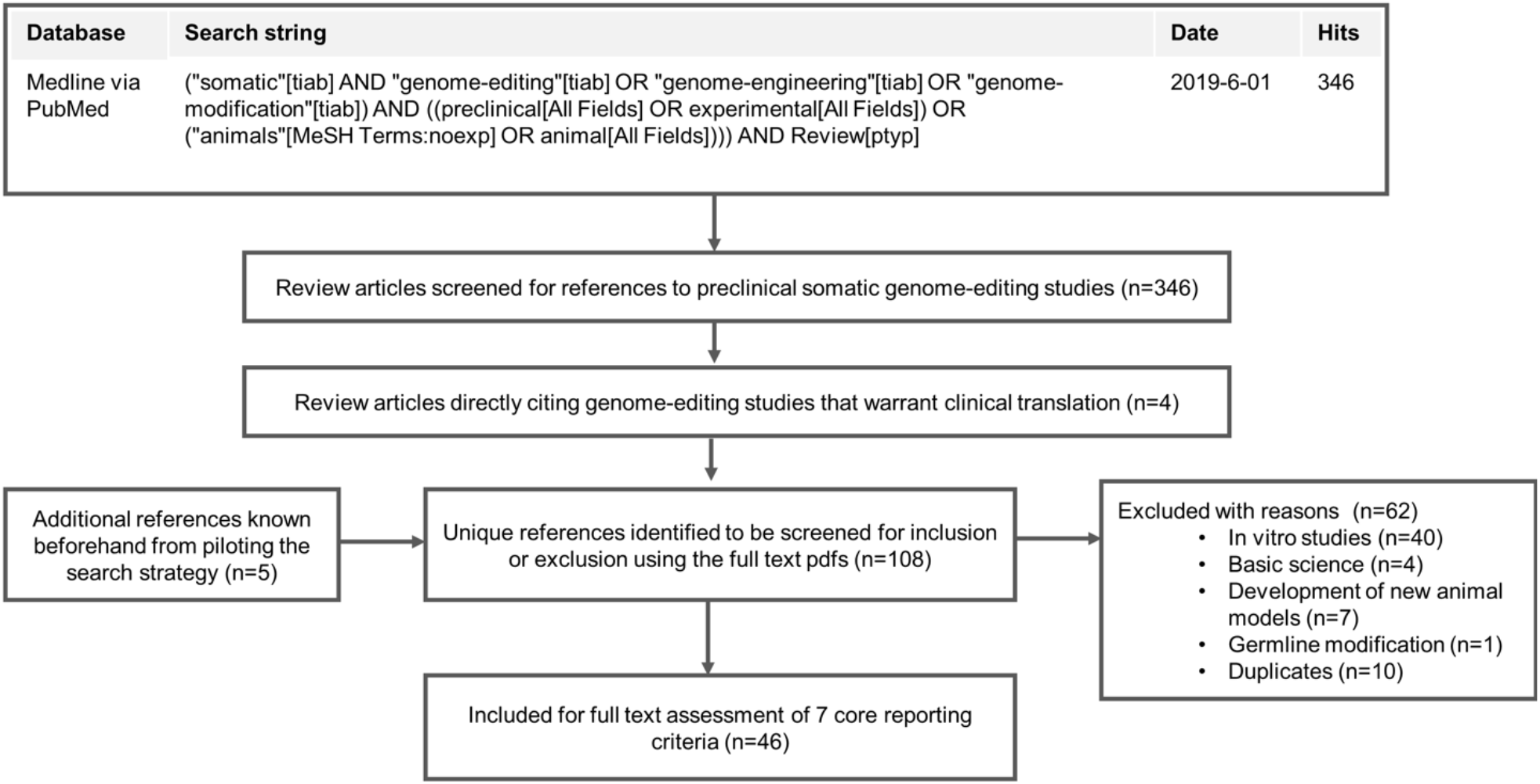
Flow diagram about the search strategy and selection process of relevant literature on preclinical research in the field of somatic cell genome editing (PubMed search link).

The reporting quality was examined by two coders (JS & MB) independently from each other using an assessment scheme in MaxQDA Pro^26^. The coding procedure tracks every decision in a fully traceable and reproducible manner. Conflicting codings were settled by discussion among all authors (JS, MB, & DS). Additional methodological details, a reproducible R script along with a list of all screened review articles and included articles can be found on the Open Science Framework (https://osf.io/bc58f/).

## Results

Our broad and sensitive search terms retrieved 346 review articles from Medline/PubMed (Figure 1). After the scoping of the search results, we identified four reviews that directly cited 46 scientific articles meeting our inclusion and exclusion criteria (https://osf.io/cf5p9/). These 46 articles were discussed in these four expert reviews as promising for clinical translation (Table 1. Most of those preclinical studies report the use of CRISPR/Cas (50%, n=23) or ZFN (35%, n=16) followed by TALEN (11%, n=5), CRISPR/Cas and TALEN (2%, n=1), or megaTal nuclease (2%, n=1) as genome editing platforms. The five most frequent therapeutic indications under investigation in the reviewed literature are Duchenne muscular dystrophy (15%, n=7), HIV infection (11%, n=5), hemophilia B (9%, n=4), and leukemia (9%, n=4) and the four journals that most frequently published the studies are Molecular Therapy (15%, n=7), Nature Biotechnology (13%, n=6), Blood (9%, n=4), and Science (9%, n=4).

**Table 1.** Absolute number and percentage of reported validity domains in preclinical research of somatic cell genome editing (SCGE) based on the full text articles.

| **Study characteristics** | **Frequency** | **Percentage (%)** |
| --- | --- | --- |
| Meeting the pre-specified inclusion criteria | 46 | 100% |
| (1) Reporting of randomization status of group allocation | 16 | 35% |
| randomly | 12 | 26% |
| non-randomly | 4 | 9% |
| (2) Reporting of blinded allocation to experimental groups | 8 | 17% |
| blinded | 3 | 6,5% |
| open-labelled | 5 | 11% |
| (3) Reporting of a sample size calculation | 7 | 15% |
| (4) Reporting the exclusion of data (animal attrition rate) | 3 | 7% |
| (5) Reporting of a pre-registered study protocol | 0 | 0 |
| (6) Multi-center study design (data collection in two labs or more) | 0 | 0 |
| (7) Reporting of independent confirmation by replication | 0 | 0 |

A minority of all assessed preclinical studies reported (1) that the group allocation of animals was performed randomly (n=12; 26%, with no study reporting the exact method of the randomization procedure), (2) that the outcome assessment was blinded (n=3; 7%), (3) how the sample size was determined (n=7; 15%), or (4) if animals or data were excluded from analysis (n=3; 7%).

Four scientific articles (9%) explicitly indicated no use of random group allocation and five articles (11%) indicated that the outcome assessment was not performed by the investigator being blinded. All other papers did not report on the methods for allocation, blinding, sample size estimation or data handling.

None of the other three measures to reduce validity threats, namely (5) pre-registration, (6) multi-centric experimental data collection, (7) independent validation, were reported in any of the studies (Table 1).

For manuscripts with accompanying supplementary material (n=41), we additionally screened the supplements for all seven items from Figure 1 (https://osf.io/bnjxr/). Information on the blinding status was reported in two supplements while the randomization status and sample size calculation were reported in one supplement, slightly increasing the total numbers shown in Figure 1.

## Discussion

Our scoping review identified 46 preclinical genome editing studies that are known in the field and discussed in expert reviews as promising for clinical translation. The majority of articles does not report on well-established measures to reduce validity threats such as randomization, blinding, sample-size calculation, and data-handling.^17,27,28^ None of these studies did report on additional, less established, but increasingly discussed and implemented measures, namely: pre-registration,^17,27,28^ multi-centric experimental data collection^20,21^, independent validation.^22^

Although the additional measures are still rare in the field of SCGE, some journals, e.g. Science, actively encourage the submission of replication studies in their editorial policy. However, the scientific merits may still be underappreciated, and replications studies may be seen as less innovative and novel. Despite such legitimate worries, replication studies are invaluable for high stakes research. In cancer research, the pharmaceutical industry has been stressing the value of independent validation for some time in order to corroborate “promising” preclinical findings and to de-risk subsequent clinical development^29,30^. As such, these novel measures are an important factor for the safe and efficient translation of preclinical findings and the bedrock to guarantee stakeholders’ confidence in preclinical studies supporting clinical translation^20,29–31^ and the practical feasibility of their implementation has already been demonstrated^32,33^.

For the interpretation of these results, some limitations warrant attention. First, we did not perform a full systematic review based on a preregistered protocol and we only included studies that were discussed in review articles as promising for clinical translation. Thus, the reviewed studies may not be representative of the larger field SCGE including purely basic science. Second, our study only assessed the current reporting practice, but the lack of reporting on the seven measures does not directly prove that the respective researchers did not employ the measures. At the moment, there are no empirical studies directly examining how many findings from preclinical studies in the field of somatic genome editing are de facto reproducible if repeated. Nonetheless, large surveys with preclinical researchers^34,35^ and the assessment of protocols of animal studies^36^ strongly suggest that these measures are in fact neglected in daily practice. This supports the interpretation that the lack of reporting also reflects the lack of application of such measures.^37^ Furthermore, the less explicitly the reporting of measures to reduce the risk of bias is, the more must readers speculate about the methods applied.

The large majority of the analyzed scientific articles are published in journals that already have a policy commending the reporting of randomization, blinding, sample size calculation, and exclusion of animals from analyses. 76.1% of the included articles were published in journals that endorse the respective reporting guideline for animal research, the ARRIVE guideline^28^, or refer to the NIH recommendations ^38^ in their editorial policies (see https://osf.io/nbuk7/). This indicates a broad acceptance of the four basic reporting criteria by journal editors and reflects the recognition of the associated scientific merits. Since the explicit reporting of these criteria will enhance the utility of scientific reports to end-users, we recommend that journal editors, peer-reviewers, and authors work closely together to strengthen the reporting of the respective methodological criteria.

The lack of reporting and use of measures to facilitate the independent critical appraisal of reported evidence is common in all areas of preclinical research.^36,39^ While all preclinical research should improve in this regard, we argue that the area of somatic genome editing has an outstanding responsibility to improve the trustworthiness and usefulness of their preclinical data. We already introduced important characteristics of the current genome editing field such as the “media hype”,^40^ the “run to the market”, and the “potential for high academic (career) rewards” in the Background section. These characteristics incentivize the quickest possible translation from pre-clinical into clinical studies.

While the speed of translational genome editing research is not inherently problematic, the important task to establish a robust benefit-risk estimation based on preclinical evidence should not be rushed. To protect patients, who voluntarily participate in clinical trials, and to deal responsibly with public funding for genome editing research, robust and reliable preclinical evidence is essential. Recent content assessment of investigator brochures (IBs) for phase I/II trials demonstrated that reported information on preclinical evidence may often be insufficient to judge a drug’s therapeutic potential.^41,42^ Furthermore, several position papers and comments from academic experts and industry representatives over the past years highlighted that the low robustness and high publication bias in preclinical research probably contributes to the high failure rates in early clinical research.^30,43,44^

Beside our arguments, two recent developments demonstrate that we are not alone in our call for a particular responsibility of genome editing research to promote the scientific quality and to develop into the lighthouse example for trustworthy and useful biomedical research.

First, the National Institutes of Health (NIH) recently modified and piloted new funding requirements for preclinical genome editing research. While the above mentioned “Landis-4” criteria^17^ were developed at the NIH and already build part of NIH funding criteria, a new additional requirement was introduced most recently for the NIH Common Fund for somatic cell genome editing.^45^ With this change, an “independent validation” of relevant preclinical studies is explicitly required before translation into phase I/II trials is warranted.^46^ Furthermore, the FDA already recommends for cellular and gene therapy products that the use of blinding or randomization “is considered” by researcher, if preclinical data is intended to serve as evidence for clinical trials.^47^

Second, in the related field of gene drive research,^37^ the implementation of an “open ethical development” to foster public discourse based on transparent open science^48^ is already acting as a lighthouse which guides the way towards open science and public preregistration. Kevin Esvelt and his MIT lab, for example, make a strong case for opening all research proposals to the public.^49^ Pre-registration of animal research receives increasing attention worldwide.^50–52^ Different to registries for clinical trials, animal study registries allows embargo times to protect against theft of ideas.^51^ Despite the embargo feature, the existing animal study registries do not reflect more than some dozens of protocols. Genome editing research could break the ice.

Our study on reporting practices supports the concern that major parts of preclinical research on genome editing do not apply measures to reduce risk of bias. While this is not specific to the field of genome editing, we argue that the responsibility to overcome this neglect is particularly high in this field. Some few lighthouse initiatives such as the NIH Common Fund^45^ and the example of Kevin Esvelt’s lab^49^ start to address this responsibility. This is the right time for societies, such as the International Society for Stem Cell Research (ISSCR), other funders and institutions to proactively join the line of first-movers. By further developing and implementing standards for “confirmation” and “pre-registration” in preclinical research the field of genome editing would become a paradigm case for trustworthy and useful translational research.

## Acknowledgements

We thank Dirk Hoffman (Hannover Medical School) for valuable feedback on a previous draft version. This work was funded by the German Federal Ministry of Education and Research (BMBF 01GP1611A, https://www.bmbf.de/). The funder had no role in study design, data collection and analysis, decision to publish, or preparation of the manuscript.

## Author contributions

**Merlin Bittlinger**: Conceptualization, Methodology, Formal Analysis, Data Curation, Writing - Original Draft, Visualization. **Johannes Schwietering**: Conceptualization, Methodology, Formal Analysis, Data Curation, Writing - Review & Editing. **Daniel Strech**: Conceptualization, Validation, Writing - Review & Editing, Supervision, Funding Acquisition.

## Data availability

All data and all R scripts underlying the results are available on the Open Science Framework (https://osf.io/bc58f/).

